# Integration of *Candida albicans*-induced single-cell gene expression data and circulatory protein concentrations reveal genetic regulators of inflammation

**DOI:** 10.1101/2022.07.25.501448

**Authors:** Collins K. Boahen, Roy Oelen, Kieu T.T Le, Mihai G. Netea, Lude Franke, Monique G.P. van der Wijst, Vinod Kumar

**Author notes:** **Corresponding author** Vinod Kumar.

## Abstract

Both gene expression and protein concentrations are regulated by genetic variants. Exploring the regulation of both eQTLs and pQTLs simultaneously in a context- and cell-type dependent manner may help to unravel mechanistic basis for genetic regulation of pQTLs. Here, we performed meta-analysis of *Candida albicans*-induced pQTLs from two population-based cohorts and intersected the results with *Candida*-induced cell-type specific expression association data (eQTL). This revealed systematic differences between the pQTLs and eQTL, with only 35% of common genetic variants modulating both intermediate molecular phenotypes. By taking advantage of the tightly co-regulated pattern of the proteins, we also identified SNPs affecting protein network upon *Candida* stimulations. Colocalization of pQTLs and eQTLs signals implicated several genomic loci including MMP-1 and AMZ1. Analysis of *Candida*-induced single cell gene expression data implicated specific cell types that exhibit significant expression QTLs upon stimulation. By highlighting the role of trans-regulatory networks in determining the abundance of proteins in blood, our study serve as a framework to gain insights into the mechanisms of genetic regulation of protein levels in a context-dependent manner.

## Introduction

Genome-wide association studies (GWAS) have successfully identified tens of thousands of associations between single nucleotide polymorphisms (SNPs) and human diseases. Correlating these GWAS SNPs with molecular traits (QTLs) such as gene expression (eQTLs) is a commonly used strategy to prioritize causal genes. This is partly due to the robustness of RNA-sequencing technologies and the feasibility of eQTLs to provide insights into the molecular mechanisms of genetic variants associated with complex diseases(Westra *et al*., 2013). The majority of the eQTL studies have made use of RNA extracted from whole blood and analyzed using bulk RNA-sequencing approaches to unravel disease biology. However, this approach is limited in identifying a genetic variant’s cell-type-specific and context-dependent impact on gene expression levels(Zhernakova *et al*., 2017; Mu *et al*., 2021) or causal cell types of a particular disease.

In addition to eQTLs, protein quantitative trait loci (pQTLs) are also important molecular traits to understand GWAS findings. Circulating plasma proteins play essential roles in various biological process such as signaling and defense against infections but also, the dysregulation of proteins themselves lead to various diseases and are mostly the targets for therapeutic interventions(Molendijk and Parker, 2021). Proteins provide the closest link to phenotypic traits as being the ultimate product of transcripts. Given that not all RNA alterations lead to functional changes, studies of genetics regulation at protein level are warranted. By making use of circulatory protein and genotype data from 30,931 samples, a recent pQTL study(Folkersen *et al*., 2020) showed that approximately 29% of the pQTLs are also GWAS SNPs. Also, pQTLs can be tissue- and context-specific. For example, SNPs affecting cytokine production upon *ex-vivo* blood stimulation were shown to overlap with SNPs associated with infectious and inflammatory diseases(Li *et al*., 2016). However, which specific cell type is contributing to the production of these circulatory proteins is unclear. In addition, by examining the relationship between pQTLs and eQTLs, previous studies have demonstrated the disparity of genetics variants associated with mRNA expressions and protein abundances as summarized by a minireview(Vitrinel *et al*., 2019), and a recently published study affirms this observation with substantial difference between pQTLs and eQTLs where only 32% of the index eQTLs variants were replicated in pQTLs(Assum *et al*., 2022). While previous studies examining the extent of overlap between pQTLs and eQTLs extensively explored steady-state conditions, the degree to which these findings are replicated in stimulated conditions remain elusive.

Proteins constitute the largest class of drug targets and thus, the identification of disease-mediating candidate proteins is crucial to bridging the gap between human diseases and the genome(Suhre, McCarthy and Schwenk, 2021). However, the proportion of pQTLs mostly overlapping with known disease-associated loci is very limited with percentages as low as 29 %(Folkersen *et al*., 2020) and 12%(Ferkingstad *et al*., 2021) observed in previous studies. Studying pQTLs in the right context could provide more explanation of the link between genetics and diseases as well potential targets for treatment. To tackle these challenges, in this study, we first aimed to identify pQTL variants upon *ex-vivo Candida albicans* stimulation in two independent European cohorts. Secondly, we combined the pQTLs identified with cell-type-specific eQTLs upon *Candida albicans* stimulation to prioritize genes at the genomic regions regulating the protein abundances. Thirdly, to examine the potential impact of these molecular or intermediate traits on a fungal sepsis, we conducted co-localization analyses with Candidemia GWAS signals. This study provides deeper insights into the genetic basis underlying variations in *Candida*-stimulated protein abundance as we identified pQTLs through meta-analysis which colocalized with cell-type-specific *cis*-eQTLs.

## Results

### Identification of protein quantitative trait loci (pQTL) in univariate manner

To identify SNPs associated with the concentration of specific proteins induced upon 24h *Candida albicans* stimulations, pQTL analysis was performed in two independent population-based cohorts. A total of 445 participants (500FG (N= 325) and 1M-scBloodNL (N=120) cohort) were studied for whom both genotype and Olink protein abundances were measured upon *ex vivo* 24h *C. Albicans* stimulation of their PBMCs (**Figure 1**). After genotype imputation and quality control, 4,095,761 SNPs and 26 inflammatory proteins remained that were common between both cohorts and that were used as input for the pQTL analysis. The protein data for both cohorts were generated in different batches. Therefore, pQTL analysis was run in each cohort separately, after which results were integrated using meta-analysis to increase statistical power. We identified a genome-wide significant *cis*-acting pQTL variant rs484915 (P value = 1.81×10^−8^) on chromosome 11 correlating with MMP-1 production (Figure 2A). In addition, the other top SNPs correlating with the remaining 25 proteins were *trans*-acting pQTLs with strong suggestive associations (P value > 5.0 ×10^−8^ to 5 ×10^−6^). For example, the second most significant hit aside the genome-wide significant cQTL, is an intronic SNP rs62205465 residing in the *ZNF133* locus and exhibited an association strength closed to the genome-wide significant threshold (P value = 5.80 ×10^−8^) with MCP-3 concentrations. **Figure 2 A** illustrates the association results of all the proteins analyzed and their corresponding top SNPs (lowest P-value).

**Figure 1:**
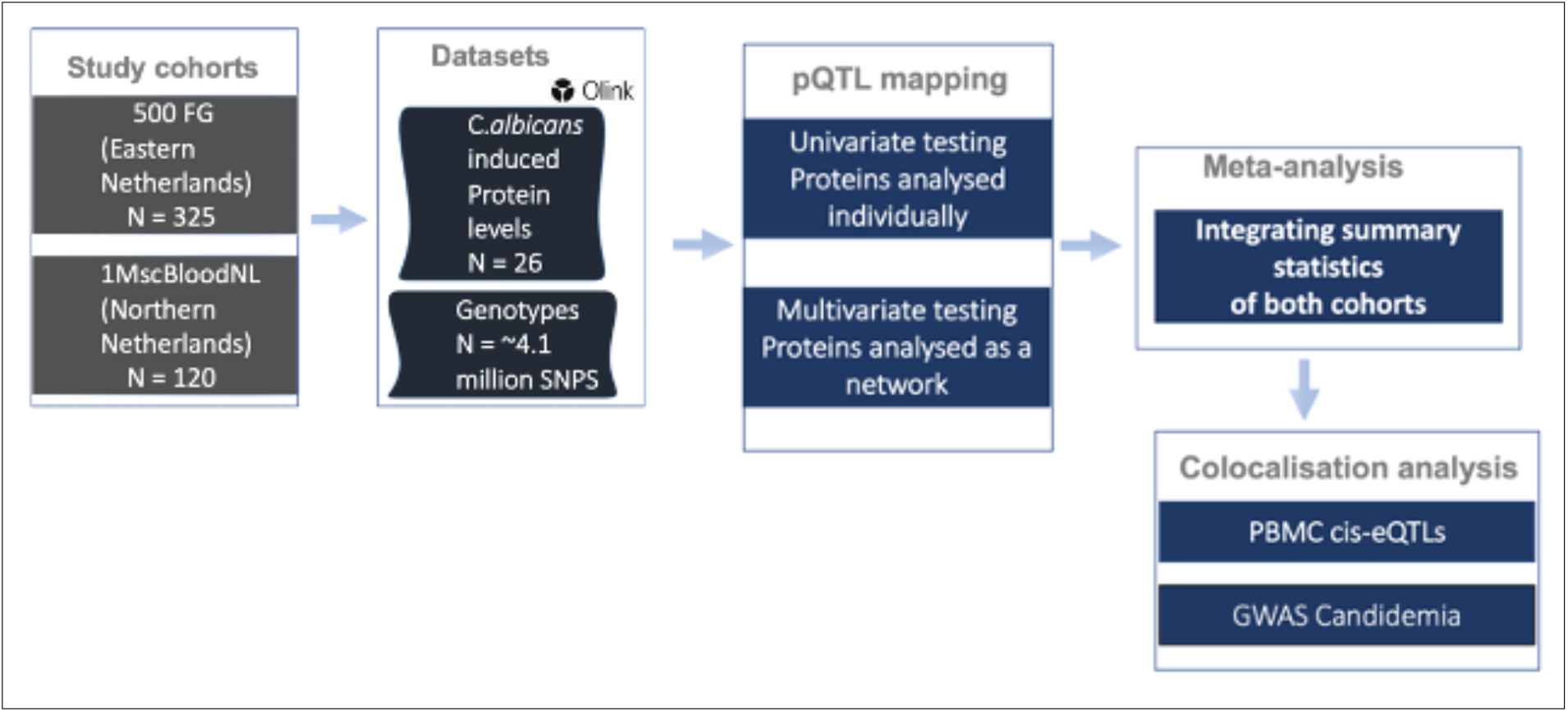
A schematic representation of study cohorts, study design and analyses conducted We performed protein quantitative trait loci (pQTL) mapping utilizing protein abundances and imputed genotypes in two population-based cohorts of individuals of European ancestry. We used two approaches to identify pQTL signals which were meta-analyzed and subsequently colocalized with cell-type specific eQTLs and candidemia GWAS signals to uncover genomic regions of shared association.

**Figure 2:**
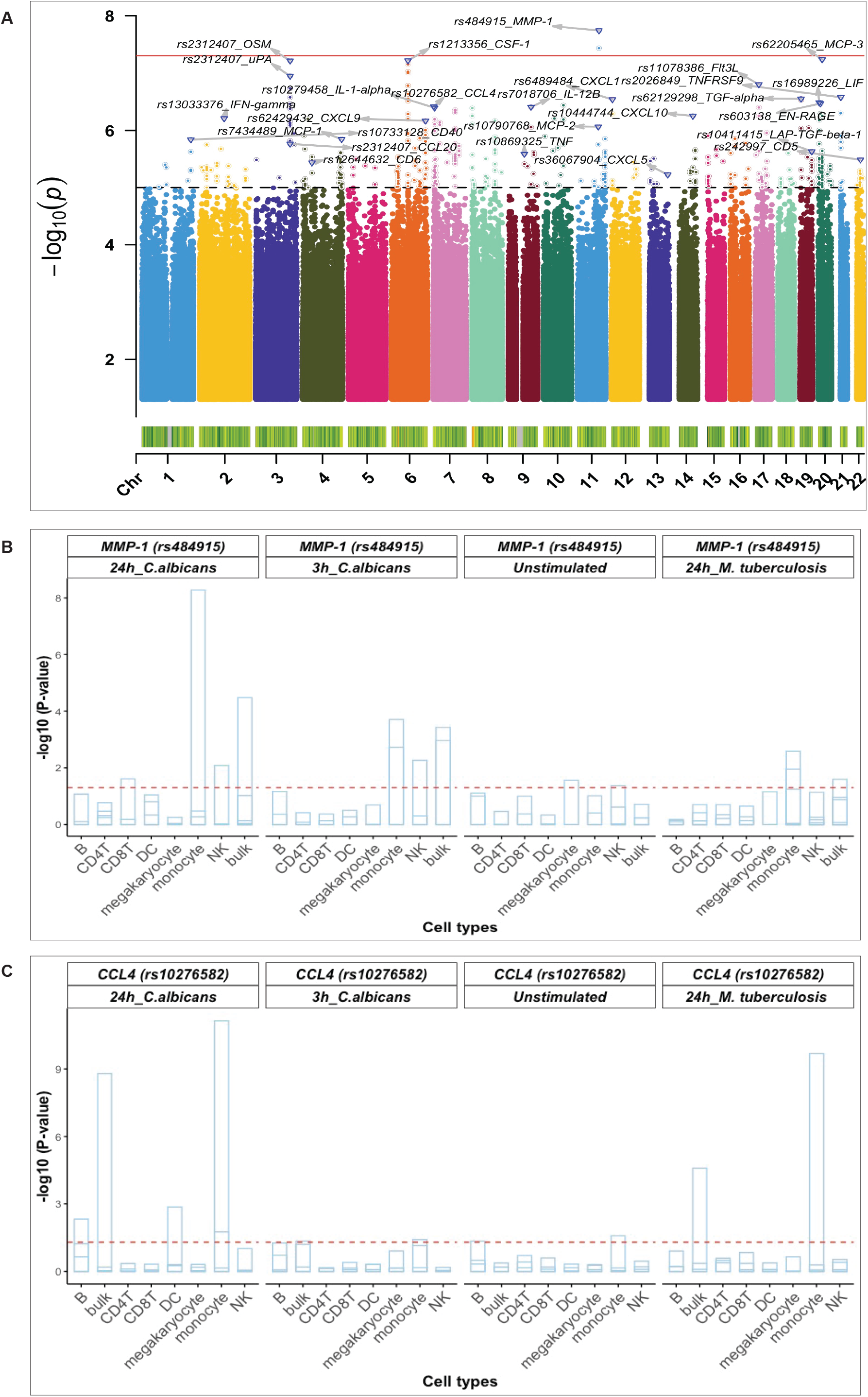
Identified pQTL SNPs and their association with cell-type specific *cis*-eQTLs (A) Manhattan plot of pQTL genetic variants associated with 26 proteins identified using a univariate approach and after meta-analysis. The red bold horizontal line depicts the genome-wide significant threshold (p-value < 1 × 10^−8^) and the black dashed denotes the suggestive association threshold (p-value = 1 ×10^−5^) Top pQTL variants and their correlated proteins are displayed on the plot. Barplots of pQTL variants correlating with - MMP-1 (B) and CCL4 (C) with genome-wide significant eQTLs in monocytes. The horizontal axis shows all the cell types considered and the vertical axis represents the negative log10 p-values for eQTLs. Each strip in the barplot corresponds to gene correlating with the pQTL variants (rs484915 and rs10276582). The barplot is also grouped per stimulation and timepoints together with pQTL variants. The horizontal red dashed line corresponds to 0.05 P-value, FDR corrected.

### Comparison between pQTL and cis-eQTL upon C. albicans stimulation

Previously, we had conducted a genome-wide eQTL analysis per major cell type using scRNA-seq data from unstimulated and 3h and 24h pathogen-stimulated (*C. albicans, M. tuberculosis, P. aeruginosa*) PBMCs in the 120 individuals from the 1M-scBloodNL cohort(Oelen *et al*., 2021). This data was used to interrogate whether the same pQTL SNP could also affect gene expression levels to explore the degree of association between pQTLs and eQTLs. Note that the pQTL data captures the bulk secretion of proteins coming from PBMCs that were stimulated for 24h with *C. albicans*, whereas the eQTL data was collected for each cell type separately. Out of the 24 unique top pQTL variants with a suggestive or genome-wide significant association (P value from 1.81 × 10^−8^ to 5.89 × 10^−6^) identified from meta-analyzed results of the univariate approach (**Table 1**), we could only overlay 20 with the scRNA-seq derived eQTL data of the 1M-scBloodNL cohort, as genotype data for the remaining 4 SNPs was not available. Seven out of the 20 tested SNPs showed an eQTL effect upon 24h *C. albicans* stimulation in at least one cell type (FDR < 0.05) (**Figure S1**). Among these seven, two pQTLs showed genome-wide significant association with gene expression levels specifically in monocytes. The first *cis*-acting pQTL, rs484915 showed association with both MMP-1 protein concentrations and gene expression (**Figure 2B**). Of note, only this pQTL among the seven was associated with the corresponding gene of the protein. The second is trans-acting SNP rs10276582 associated with CCL4 protein production affected *AMZ1* gene expression levels (**Figure 2C**). This observation suggests *AMZ1* may regulate CCL4 protein levels depending on an individual’s genotype at SNP rs10276582 (or any other SNP in high LD).

**Table 1:**
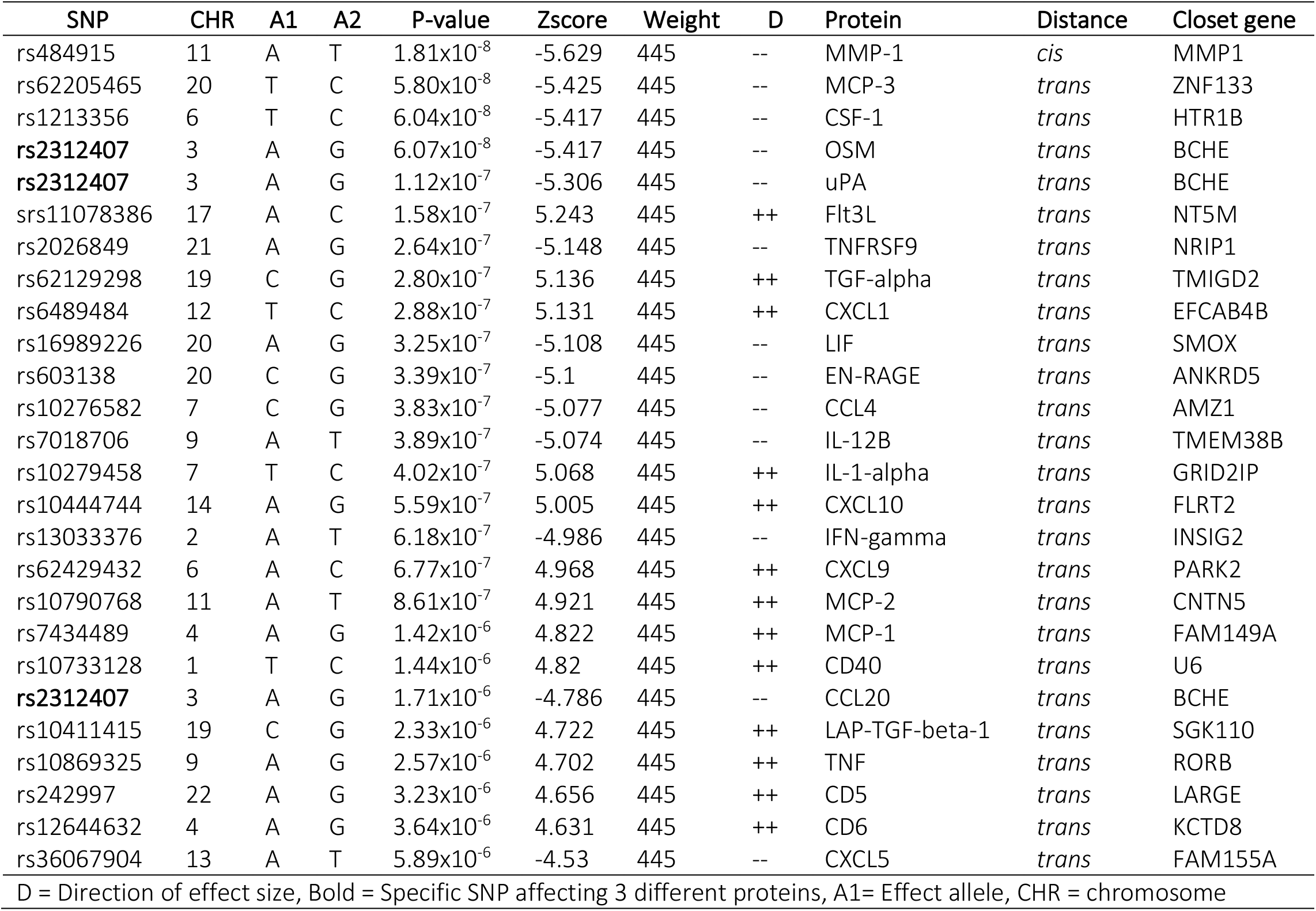
Summary of the top pQTL from meta-analysis of 26 proteins common in 500FG and 1M-scBloodNL

| SNP | CHR | A1 | A2 | P-value | Zscore | Weight | D | Protein | Distance | Closet gene |
| --- | --- | --- | --- | --- | --- | --- | --- | --- | --- | --- |
| rs484915 | 11 | A | T | $1.81 \times 10^{-8}$ | -5.629 | 445 | -- | MMP-1 | *cis* | MMP1 |
| rs62205465 | 20 | T | C | $5.80 \times 10^{-8}$ | -5.425 | 445 | -- | MCP-3 | *trans* | ZNF133 |
| rs1213356 | 6 | T | C | $6.04 \times 10^{-8}$ | -5.417 | 445 | -- | CSF-1 | *trans* | HTR1B |
| **rs2312407** | 3 | A | G | $6.07 \times 10^{-8}$ | -5.417 | 445 | -- | OSM | *trans* | BCHE |
| **rs2312407** | 3 | A | G | $1.12 \times 10^{-7}$ | -5.306 | 445 | -- | uPA | *trans* | BCHE |
| srs11078386 | 17 | A | C | $1.58 \times 10^{-7}$ | 5.243 | 445 | ++ | Flt3L | *trans* | NT5M |
| rs2026849 | 21 | A | G | $2.64 \times 10^{-7}$ | -5.148 | 445 | -- | TNFRSF9 | *trans* | NRIP1 |
| rs62129298 | 19 | C | G | $2.80 \times 10^{-7}$ | 5.136 | 445 | ++ | TGF-alpha | *trans* | TMIGD2 |
| rs6489484 | 12 | T | C | $2.88 \times 10^{-7}$ | 5.131 | 445 | ++ | CXCL1 | *trans* | EFCAB4B |
| rs16989226 | 20 | A | G | $3.25 \times 10^{-7}$ | -5.108 | 445 | -- | LIF | *trans* | SMOX |
| rs603138 | 20 | C | G | $3.39 \times 10^{-7}$ | -5.1 | 445 | -- | EN-RAGE | *trans* | ANKRD5 |
| rs10276582 | 7 | C | G | $3.83 \times 10^{-7}$ | -5.077 | 445 | -- | CCL4 | *trans* | AMZ1 |
| rs7018706 | 9 | A | T | $3.89 \times 10^{-7}$ | -5.074 | 445 | -- | IL-12B | *trans* | TMEM38B |
| rs10279458 | 7 | T | C | $4.02 \times 10^{-7}$ | 5.068 | 445 | ++ | IL-1-alpha | *trans* | GRID2IP |
| rs10444744 | 14 | A | G | $5.59 \times 10^{-7}$ | 5.005 | 445 | ++ | CXCL10 | *trans* | FLRT2 |
| rs13033376 | 2 | A | T | $6.18 \times 10^{-7}$ | -4.986 | 445 | -- | IFN-gamma | *trans* | INSIG2 |
| rs62429432 | 6 | A | C | $6.77 \times 10^{-7}$ | 4.968 | 445 | ++ | CXCL9 | *trans* | PARK2 |
| rs10790768 | 11 | A | T | $8.61 \times 10^{-7}$ | 4.921 | 445 | ++ | MCP-2 | *trans* | CNTN5 |
| rs7434489 | 4 | A | G | $1.42 \times 10^{-6}$ | 4.822 | 445 | ++ | MCP-1 | *trans* | FAM149A |
| rs10733128 | 1 | T | C | $1.44 \times 10^{-6}$ | 4.82 | 445 | ++ | CD40 | *trans* | U6 |
| **rs2312407** | 3 | A | G | $1.71 \times 10^{-6}$ | -4.786 | 445 | -- | CCL20 | *trans* | BCHE |
| rs10411415 | 19 | C | G | $2.33 \times 10^{-6}$ | 4.722 | 445 | ++ | LAP-TGF-beta-1 | *trans* | SGK110 |
| rs10869325 | 9 | A | G | $2.57 \times 10^{-6}$ | 4.702 | 445 | ++ | TNF | *trans* | RORB |
| rs242997 | 22 | A | G | $3.23 \times 10^{-6}$ | 4.656 | 445 | ++ | CD5 | *trans* | LARGE |
| rs12644632 | 4 | A | G | $3.64 \times 10^{-6}$ | 4.631 | 445 | ++ | CD6 | *trans* | KCTD8 |
| rs36067904 | 13 | A | T | $5.89 \times 10^{-6}$ | -4.53 | 445 | -- | CXCL5 | *trans* | FAM155A |

D = Direction of effect size, Bold = Specific SNP affecting 3 different proteins, A1= Effect allele, CHR = chromosome

To elucidate whether the effects of the significant eQTLs were detectable only in *Candida* stimulations after 24h, we further assessed their impact in unstimulated condition, 3h *Candida* stimulations, as well as in *Mycobacterium tuberculosis* stimulations after 24h. We observed that the effects of the *cis*-eQTLs were less pronounced before stimulation and after 3h *Candida* stimulation, which suggests temporal regulation of gene expression following *Candida* stimulation. Nearly equal strength of association was identified in the case of *M. tuberculosis* stimulations after 24h for only the *trans*-acting variant SNP rs10276582 (**Figure 2C**).

### Colocalization analysis identifies causal genes at C. albicans-induced pQTLs

Next, to uncover the potential mechanisms underlying the observed pQTLs, we tested whether SNPs impacting protein concentrations at specific loci are also the same causal regulatory eQTLs through colocalization analysis. We identified strong evidence of colocalization for both pQTLs and eQTLs. For instance, in the MMP-1 locus (**Figure 3A**), the posterior probability (PP.H4) was 0.998 (**Figure 3B**). Also, the *trans*-acting pQTL SNP rs10276582 located in the *AMZ1* locus on chromosome 7 (**Figure 3C**), showed significant colocalization (PP.H4 = 0.996) with the eQTL effect. eQTL analysis revealed association of this variant with three different *cis* genes, *GNA12* (P = 7.1 × 10^−1^), *TTYH3* (P = 1.7 × 10^−2^) and *AMZ1* (P = 7.31 × 10^−12^) (**Figure 3D**), indicating the presence of multiple causal genes at this locus. However, both *GNA12* and *TTYH3* exhibited weaker strength of association than AMZ1, which showed statistically significant association. Thus, pinpointing AMZ1 as the potential causal gene in this genomic region. This observation demonstrates that *cis*-regulation of gene expression levels maybe involved in the mechanisms by which distal variants impact protein expression.

**Figure 3:**
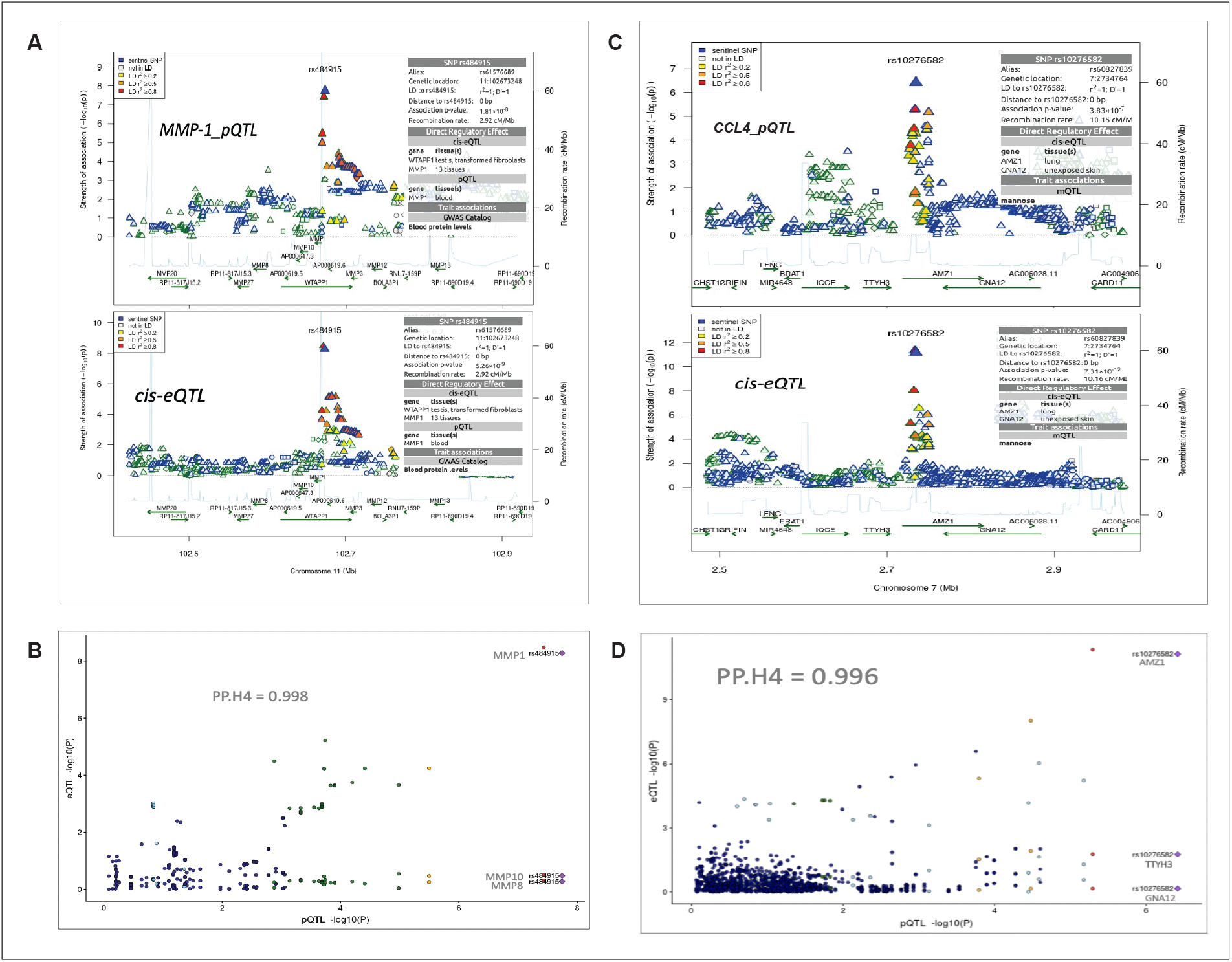
Summary of colocalization analyses between pQTL and eQTL (A) Regional association plots at the MMP-1 locus for MMP-1 protein QTL result (top panel) and cis-QTL expressions result (bottom panel) on chromosome 11. (D) Regional association plots at the AMZ1 locus with CCL4 protein QTL result (top panel) and cis-QTL expressions result (bottom panel) on chromosome 7. The sentinel SNPs (rs484915 and rs10276582) are indicated with blue diamond shape and other surrounding SNPs are colored with different levels of linkage disequilibrium with the sentinel SNPs. The horizontal axis indicates chromosomal positions (NCBI human genome build 37) and the vertical axes represent negative log_10_ p-values and recombination rates (cM/Mb) estimated from 1000 Genome Project (European population) version 3.3. (B and D) Correlation plots with strong evidence of colocalization between pQTLs and eQTLs as indicated by PP.H4 (posterior probability of shared causal variants) values.

#### Exploring other mechanisms of pQTLs function on protein levels

To further understand the mechanisms or regulation of the trans-pQTL, we first looked for evidence of the genes near or at the trans loci encoding for any of the proteins interacting with our tested protein. To do this, we used an annotation database – Human Integrated Protein-Protein Interaction Reference (HIPPIE)(Alanis-Lobato, Andrade-Navarro and Schaefer, 2017) and defined trans genes as all genes falling within 1Mb window centered on the top 25 identified trans-pQTL loci. Generally, we did not observe any interacting partners between the trans genes and the trans affected proteins (**Figures S2-3**).

### Co-regulation of C. albicans-induced protein levels

It is possible that some of the genetic variants may affect multiple proteins, so we wanted to explore whether there is strong correlation between protein concentrations upon stimulation. To explore the patterns underlying *Candida*-induced protein production, we performed correlation analyses. We observed mostly significant positive pairwise correlation (with the exception of MCP-1 and MCP-2) between the 26 proteins in the 500FG cohort, which contains the largest number of individuals from which samples were collected (**Figure 4A**). However, in the 1M-scBloodNL cohort, divergent patterns of correlation strengths were observed, including weaker and negative correlations between CSF-1, IL-12B and IFN-gamma, contrasting the positive correlation observed in the 500FG cohort (**Figure 4B**). To understand what might underlie this observation, we performed the Fligner-Killen test to evaluate whether there is significant donor variation of these proteins between the two cohorts. While we observe significant difference for CSF-1 (Fligner-Killeen:med chi-squared = 19.893, p-value = 8.19 × 10^−6^) and IL-12B (Fligner-Killeen:med chi-squared = 5.0642, p-value = 0.02442), there was no evidence to suggest that the variance in IFN-gamma concentrations significantly differ between both cohorts (Fligner-Killeen:med chi-squared = 0.3097, p-value = 0.5779).

**Figure 4:**
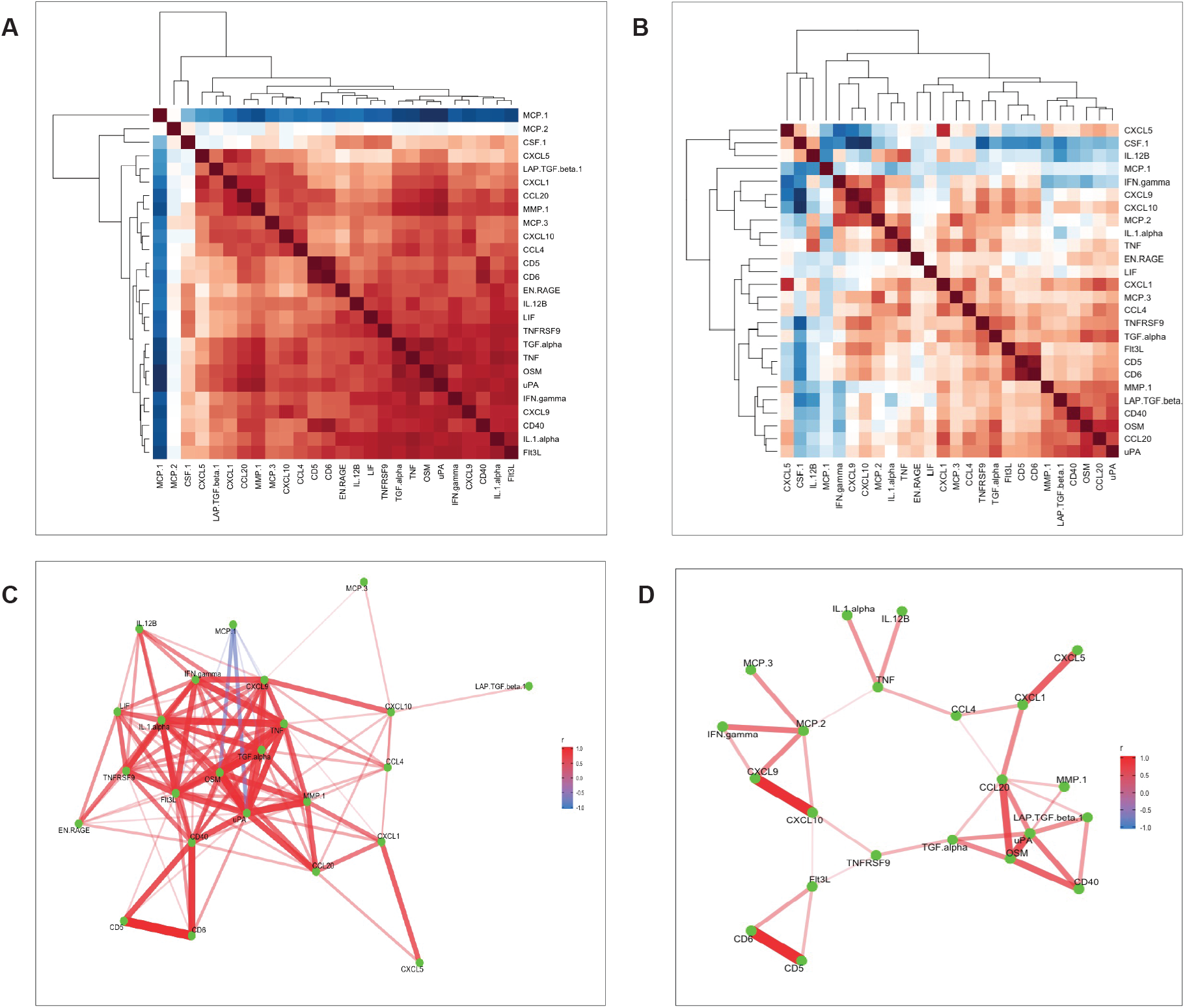
Heatmaps of correlation analysis (A and B) Correlation of 26 common proteins for the 500FG and 1MscBloodNL cohorts respectively. Pairwise correlation between proteins were computed using residual values after adjusting age and sex on protein levels. (C and D) Represents network of proteins with strong pairwise correlation coefficients for the 500FG and 1M-scBloodNL cohorts respectively.

In the 500FG cohort, the strongly pairwise correlated proteins acting as protein network consisted of 24 out of the 26 proteins common between both cohorts (**Figure 4C**). For example, the correlation coefficient between uPA and OSM was as high as 0.91, while CD40 exhibited an approximately 0.9 between CD5 and CD6. Next, we sought to more accurately infer the protein network by replicating the analysis in the 1M-scBloodNL cohort. While MCP-1 exhibited strong negative correlation with some proteins (such as OSM and uPA) in the 500FG cohort, this pattern of correlation was hidden in the network obtained from the 1M-scBloodNL cohort (**Figure 4D**). A rather weaker and positive correlation was manifested between MCP-1 and OSM (0.26) and uPA (0.24) and thus, MCP-1 was excluded from multivariate genetic association analysis as shared genetic architecture is likely to underlie or trigger the observed strong positive correlations.

### Genome-wide identification of pQTLs impacting protein network

Taking advantage of the tightly co-regulated pattern of the proteins, we sought to identify SNPs impacting protein network upon *Candida* stimulations aside the univariate association analysis. After independent analyses in both cohorts, we identified a genome-wide significant hit in the 500FG cohort as well as several suggestive associations. The genome-wide significant SNP was rs938662 (P value = 4.37 × 10^−8^), residing in the *PRKCE* locus (**Figure S4A**). There was no evidence of inflation of the test statistics as genomic inflation factor (*λ*) was computed to be 1.01 (**Figure S4B**). In the case of the 1M-scBloodNL dataset, no statistically significant SNP was found to be associated with proteins (**Figure S4C**), and corresponding genomic inflation factor (*λ*) was 0.99, indicating lack of inflation of test statistics (**Figure S4D**). To identify true or robust association signals, we synthesized summary statistics from both cohorts through meta-analysis. Even though no loci reached genome-wide significance, we identified strong suggestive associations (**Figure 5A**). The top signal identified was rs484915 (P value = 1.05×10^−7^), located at the *MMP1* locus and has previously been shown to alter expression levels of *cis*-genes in multiple tissues and also blood protein concentrations(Suhre *et al*., 2017). In the univariate analysis, we found the same SNP rs484915 to be associated with MMP-1 with slightly much stronger association based on P-value (1.81×10^−8^). Interestingly, five proteins, namely MMP-1, CXCL5, CCL20, CXCL1 and CXCL10 among the proteins forming the network (**Figure 5B**), contributed with relatively stronger weights or correlation coefficients to the observed association result of SNP rs484915 to the protein network. This observation from the multivariate approach demonstrates the pleiotropic effect of SNP rs484915 which cannot be captured directly via univariate analysis and also tease apart the main proteins whose expression levels are being regulated.

**Figure 5:**
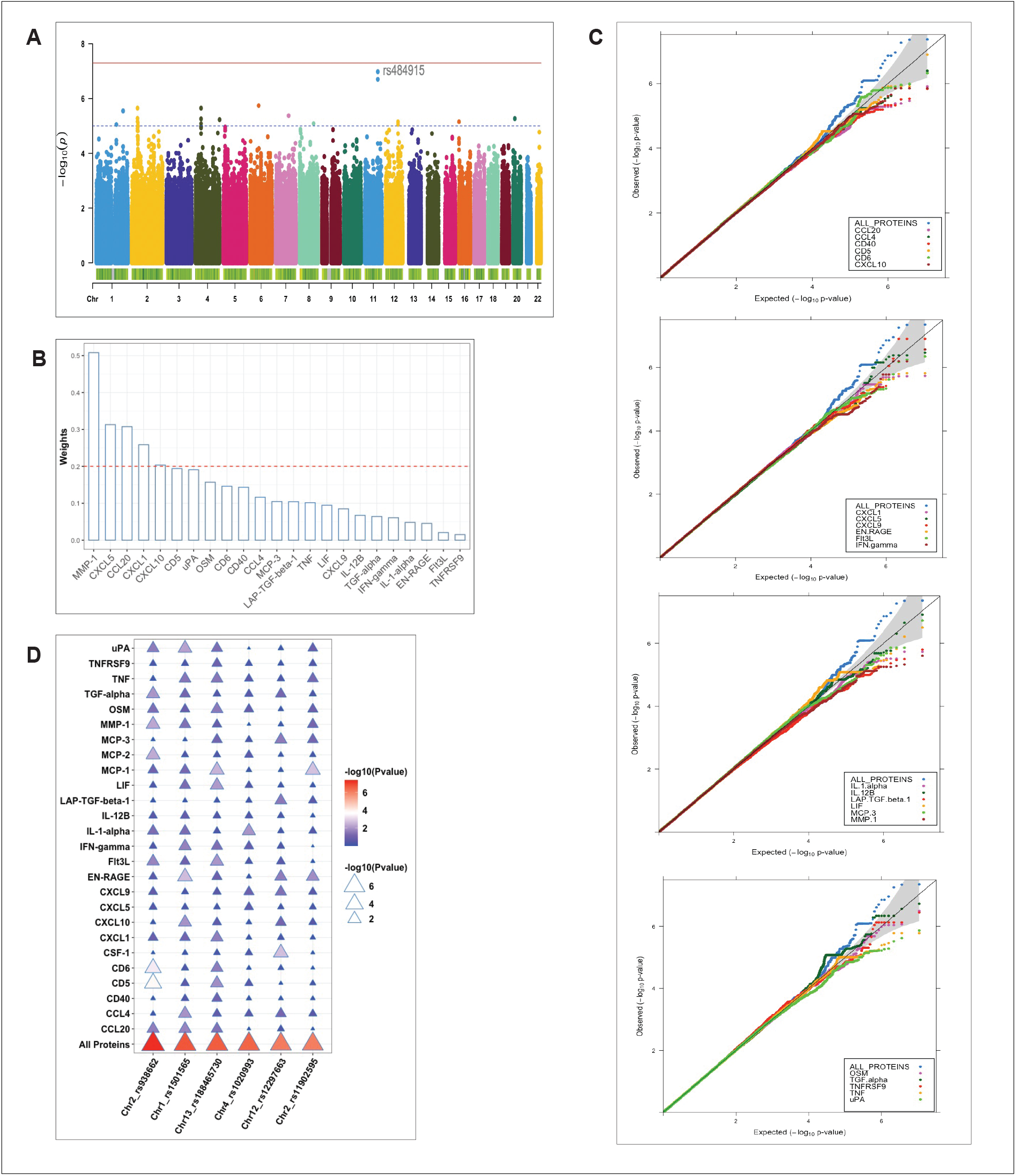
Summary of multivariate QTL mapping results and comparison with pQTLs identified using univariate approach (A) Manhattan plot for meta-analyzed pQTL multivariate analysis results of protein network. Strength of association (-log10 meta-p values) is shown on the vertical axis and the horizontal axis depicts the chromosomal position of plotted SNPs. The blue horizontal dashed line and the red bold line represent suggestive (p-value = 1 ×10^−5^) and genome-wide (p-value < 5 ×10^−8^) significant thresholds, respectively. (B) Barplot of the weights (contribution to the protein network association) on the vertical axis plotted against all proteins forming the network (horizontal axis) using “ggbarplot” function in R. The red dashed line represents the threshold of significant contributions. (C) Quantile-quantile (Q-Q) plots for the pQTL mapping results in the 500GF cohort. The p-values distribution of the multivariate analysis (ALL PROTEINS) results are shown in blue and the remaining colors correspond to p-values of proteins analyzed separately. The gray shaded area represents 95% confidence interval of the null hypothesis. (D) Plot of association results of all individual proteins and protein network (All Proteins) plotted against the top six independent SNPs (horizontal axis) identified after multivariate analysis in the 500FG cohort. The color legend represents strength of association of SNPs with protein levels (Pvalue), ranging from blue (weaker associations) to red (stronger associations).

### Multivariate approach improves statistical power

Next, we sought to investigate whether the joint analysis of multiple correlated proteins with genetic variants offers some advantage over univariate analysis. To achieve this aim, we directly compared the P-values or distribution of P-values of genetic variations identified using both approaches and restricted the analysis to only the largest cohort, 500FG. We reasoned that such analysis will be more credible to be conducted in a specific cohort due to the variation seen in proteins’ correlation structure between both cohorts. As expected, we observed stronger associations emanating from the multivariate analysis compared to the univariate manner (**Figure 5C**). The top 6 independent strong suggestive loci (P < 9 ×10^−7^) identified via the multivariate approach were in all cases showing a much stronger association when compared to the strength of association (P values) of each protein analyzed separately (**Figure 5D**). Furthermore, to characterize the identified pQTLs, we targeted the top 6 strongest pQTL variants, given the lack statistically significant SNPs to elucidate their effect on cell-type specific eQTLs. We found 3 out of 6 to be an eQTL as well in at least one cell type albeit with nominally significant effect. The overlapped SNPs showed weak association with various *cis*-genes. For instance, the intronic SNP rs11902595 on chromosome 2 was nominally (P= 0.013) correlated with a lincRNA (RP11-191L7.1) in CD8T cells.

### Overlap of pQTLs with SCALLOP consortium data

We further evaluated the overlap and strength of association between our *Candida* 24h stimulated pQTL associations and previously reported highly powered (21,758 participants) unstimulated pQTL study published under the SCALLOP consortium(Folkersen *et al*., 2020). We found only 8 proteins (MMP-1, CSF-1, CXCL1, EN-RAGE, CCL4, MCP-1, CD40, CCL20) common between the 90 cardiovascular proteins measured in the previous study and the 26 proteins measured in the inflammatory panel used in our study. Using nominal significance P-value < 0.05, the percentage of shared *Candida* 24h stimulated pQTLs vs pQTLs identified using the SCALLOP consortium data, ranged from 4.5% to 5.6 %. Of these, the top SNPs among the shared variants show nominal association with 6 proteins after stimulation. However, *cis*-acting genome-wide significant pQTL variant rs484915 (P = 1.81 × 10^−8^) correlating with MMP-1 upon stimulation showed very strong association (P = 1.87 × 10^−220^) in the SCALLOP consortium data (**Figure 6A**). We further observed one *trans*-acting pQTL variant rs3014874 exhibiting suggestive association with EN-RAGE protein (5.87 × 10^−4^), but exceeded the genome-wide significant threshold with P = 1.64 × 10^−29^ in the SCALLOP consortium data (**Figure 6B**). Evidence from previous studies indicate that this downstream variant (rs3014874) located on chromosome 1 affects the expression levels of *cis*-genes in blood as well as different tissues (**Figure 6C**). Given that eQTL effects are dependent on context and tissue being studied, we investigated the effect on this SNP on nearby genes in different cell types after candida stimulation. Among all the cis-genes, SNP rs3014874 showed association with only *S100A9* gene (**Figure 6D**), making it the likely causal gene. This observation further highlights the role of trans-regulatory network in determining the abundance of circulating plasma proteins in blood.

**Figure 6:**
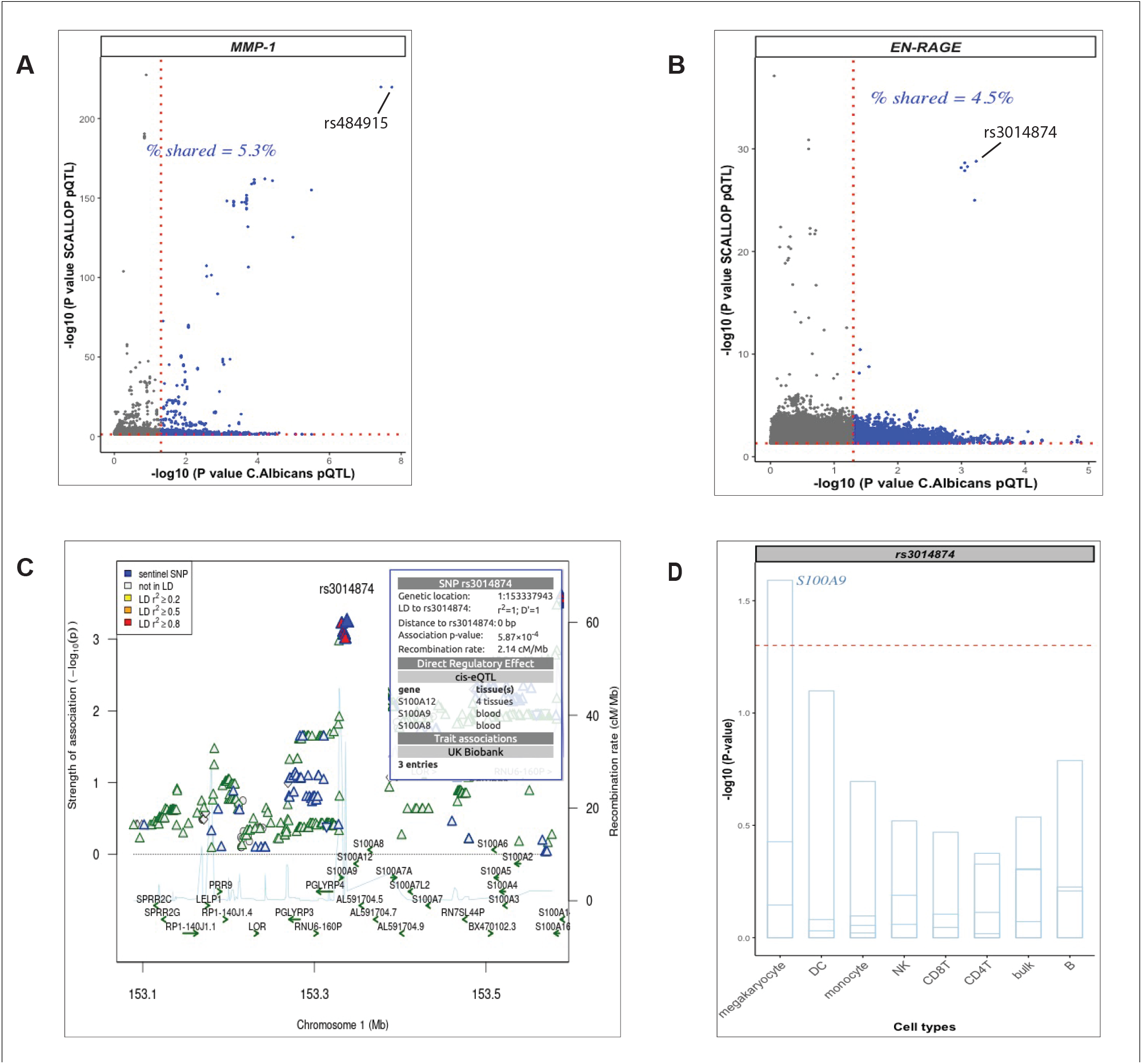
Exploration of pQTL variants in plasma from unstimulated samples (A and B) Scatter plots generated with the “ggbarplot” function of pQTLs from unstimulated protein levels (SCALLOP) against candida-induced protein levels showing the proportion of shared pQTLs. Top cis-acting pQTL and trans-acting pQTL are labelled respectively. Dot gray color represents unique pQTLs from the SCALLOP data analysis without stimulation and dot blue color represents the number of overlapping pQTLs irrespective of stimulation status. (C) Regional association plot at the S100A9 locus using EN-RAGE pQTL association results after candida stimulation. The top SNP rs3014874 on chromosome 1, with direct regulatory effect on multiple cis-genes is indicated in blue diamond shape. (D) Barplot of the top trans-acting pQTL (rs3014874) association on multiple cis-genes (strips in the barplot) in the S100A9 locus in various cell types (horizontal axis). At nominal level (red dashed horizontal line), SNP rs3014874 significantly affect only S100A9 gene in megakaryocyte.

### Limited evidence of pQTL signals colocalizing with candidemia GWAS susceptibility signals

We further sought to identify candidemia loci sharing a genetic signal with pQTLs to uncover the link between pQTLs and candidemia susceptibility. Among the 8 genomic loci investigated (**Table S1**) using data from previously published GWAS(Jaeger *et al*., 2019), only two genomic regions showed weak evidence of a shared single causal variant with multiple proteins. In the *EHD4* locus on chromosome 15 for instance, PP.H4 was estimated to be 0.22 and 0.16 for urokinase-type plasminogen activator (uPA) and Flt-3 ligand (Flt3l) respectively (**Figure S5A**). Similarly, at the *POU4F1* locus, even though SNP rs6563046 was the index variant for both pQTL and GWAS analyses, the estimated PP.H4 for CCL4 and CXCL1 were approximately as low as 0.10 (**Figure S5B**). This observation suggests that SNPs affecting excreted proteins may have limited role in candidemia.

## Discussion

In this study we applied integrative analysis of genomics, proteomics and single cell transcriptomics in two independent cohorts to better understand the genetic mechanisms that link mRNA expression and proteins abundance following *Candida albicans* stimulation of immune cells. First, we compared univariate versus multivariate pQTL analyses to test how the cross-trait covariance information which is mostly unutilized in the univariate analysis influence pQTL findings. We then overlaid the identified pQTL SNPs with cell-type-specific eQTL results from PBMCs to help disentangle the underlying mechanism of pQTL results and specify the cell type that could be involved. Finally, we determined whether the identified pQTLs were colocalizing with GWAS SNPs for candidemia susceptibility, providing us with the potential role of secreted proteins in the genetic susceptibility for candidemia. The strength of this study is the meta-analytic approach of combining two independent population-based cohorts which makes it possible to identify true or consistent genetic associations. Also, for the genes involved in many phenotypes or complex diseases’ progression, it is unclear in which cell type gene regulation takes place. Thus, the application of cell-type-specific *cis*-eQTL data addresses this challenge.

One of the main observations from our study is that only 35% (7/20) of the pQTLs significantly correlated with mRNA expressions at single cell level, suggesting that eQTLs cannot be used as a proxy for pQTLs when investigating molecular mechanisms underlying trait-associated variants. The observation of limited overlap is consistent with previously reported findings as the discrepancy between pQTL and eQTL results were also detected in a larger cohort (GTEx Consortium) utilizing over 900 individuals(Carithers and Moore, 2015). Another recent proteomic study also demonstrated that more than 2000 protein associated variants had no eQTLs(He *et al*., 2020). Of note, in each of those studies the protein data was a bulk measurement from circulating proteins in the blood (potentially being secreted by any cell type in the body), whereas the matched mRNA data was a bulk measurement from the immune cells themselves. Similarly, in our own study the protein data was a bulk measurement from PBMCs, whereas scRNA-seq data was used to obtain cell-type-specific eQTL data. This discrepancy can potentially result in different significant/top SNPs being identified in the bulk pQTL versus the cell-type-specific eQTL analysis. Given the recent emergence of high-throughput technologies(Chung *et al*., 2021; Reimegård *et al*., 2021) to measure both mRNA and protein levels simultaneously at single cell resolution, future studies could directly compare cell-type-specific pQTL and eQTL results from exactly the same samples. This will provide the most definitive answer regarding the eluded low overlap of both data modalities. Even so, this finding suggests that protein regulation is much more complex than direct mRNA-protein relationship: this is not necessarily surprising, as many post-transcriptional processes are known to influence protein production such as translation, processing and secretion. Therefore, on average low correlation between mRNA and protein QTLs was not unexpected. For instance, 6 different regulatory patterns have been previously described as the mechanisms by which genetic variants can influence the process of transcription to translation(Wang *et al*., 2019), such as in scenarios whereby SNPs independently affect transcript levels and protein abundance or SNPs responsible for both transcriptional and translational alterations. Thus, we advocate for future studies targeting genome-wide *Candida albicans*-induced mRNA (*cis* and *trans* inclusive) and cell-type-specific protein expressions analysis which has the potential to refine this observation and makes it feasible for genome-wide comparison of the proportion of shared or unique pQTL and eQTL variants.

To help with the interpretation of how cell-type-specific eQTLs regulate trans-acting pQTLs, colocalization analysis of lead pQTLs and eQTL signals implicated the *AMZ1* gene. This finding implies that *AMZ1* gene might be involved in the molecular pathways underlying complex diseases, most probably the pathogenesis of candidemia. Also, our analysis therefore predicts the sentinel SNP rs1027658 associated with the trans-acting protein (CCL4) as a probable causal variant and further shows that regulation of CCL4 protein levels is mediated by gene transcription. Apart from the trans-genomic region, similar analysis also showed strong colocalization at the *MMP1* locus with the leading SNP (rs484915) located in the *cis* region, suggesting a direct regulatory effect on MMP-1 protein concentrations.

Another interesting observation made in this study is the strong correlation among proteins concentrations released by human PBMCs upon *Candida albicans* stimulation, suggesting their concerted role in immune regulation. In genetic studies, joint analysis of correlated phenotypes in a single model, a so-called multivariate approach, has been demonstrated to increase statistical power relative to a univariate approach(Inouye *et al*., 2012; Yang and Wang, 2012; Zhou and Stephens, 2014). Indeed, this was clearly demonstrated in the larger cohort (500FG). For example, as the intergenic SNP rs938662 was statistically significant when the multivariate method was adopted, the lowest P value of the same SNP in the univariate approach showed suggestive association (1.45 ×10^−4^), correlating with CD5 proteins levels and strikingly, did not show any association with as many as 16/26 proteins analyzed separately.

However, the added value of coupling proteins for joint genetic analysis was not substantially detected after meta-analysis of both cohorts as we expected since multivariate analysis is known to perform relatively better especially, in the case of presence of pleiotropy(Zhang *et al*., 2009). The dissimilarity in the correlation structure between the two datasets and the relatively weaker correlation strength in the 1M-scBloodNL cohort is mostly capable of causing the multivariate test from not out performing or boosting power than the univariate test after the combined analysis. Even though smaller sample sizes can lead to instability in estimating correlation coefficient(Sari *et al*., 2017), technical and experimental variations can partially explain the observed differences in correlation pattern seen in both datasets. Yet, findings from the largest cohort demonstrates that these two approaches are entirely orthogonal to detecting genotype-phenotype relationship.

To further explore the functional consequences of the genetic variants identified, we also utilized pQTLs to understand the function of genetic loci identified in a candidemia GWAS through colocalization analysis. Even though some top SNPs were common for both traits in certain genomic regions, colocalization analysis did not implicate any strong shared genetic loci between pQTLs and GWAS signals. Given that the GWAS data is not well-powered and without statistically significant loci, it may be premature to conclude that pQTLs do not mediate the link between genetic variants and candidemia disease. Indeed, genetic variants appearing to be shared at a specific locus between pQTL and GWAS signals could be driven by linkage disequilibrium and not necessarily shared causal variants. Larger-scale candidemia GWAS studies would be useful to disentangle these possibilities. Supporting this argument, a recent powered study evaluating the relationship between pQTLs and GWAS loci of 81 diseases and other clinical traits found 69 out of 76 (number of phenotypes associated with the genome-wide significant loci investigated) representing 90.8% of the tested genetic associations with phenotypes were also associated with at least one protein with strong evidence of colocalization(Gudjonsson *et al*., 2022).

Several limitations of the present study are worth highlighting. First, the use of a specific Olink panel with overrepresentation of inflammatory proteins hinders broad analysis of proteins as the current high-throughput Olink Explore panel is capable of profiling thousands of plasma proteins. Second, sample size limitation made it impossible to comprehensively characterize pQTLs. We therefore acknowledge that upscaling the sample size might help identify more significant genetic loci especially distal QTLs with relatively smaller effective sizes and thus requiring large sample sizes to be detected.

In conclusion, our study has pinpointed several possible mechanisms through which protein expressions are regulated and also delineate the specific cell type involved. In addition, we have prioritized candidate genes at pQTL loci, providing great insight into the genetic architecture of protein levels following *C. albicans* stimulations.

## Materials and methods

### Study populations

The 500FG cohort of healthy individuals of Western European ancestry comprises of 237 males and 296 females with age range of 18 to 75 years, being part of the Human Functional Genomics Project, HFGP (www.humanfunctionalgenomics.org).

The 1M-scBloodNL cohort comprises of 120 individuals, 53 males and 67 females between 27 to 78 years. This cohort is part of the Lifelines DEEP cohort, a prospective population cohort of participants from the northern Netherlands(Tigchelaar *et al*., 2015).

### PBMC isolation and Candida albicans stimulation experiments

#### 500FG cohort

Peripheral blood mononuclear cells (PBMCs) collection has been previously described(Li *et al*., 2016). With informed consent, venous blood was drawn from the cubital vein of study participants into 10mL EDTA Monoject tubes (Medtronic, Dublin). The fraction of PBMC was obtained by density centrifugation of EDTA blood diluted 1:1 in pyrogen-free saline over Ficoll-Paque (Pharmacia Biotech,Uppsala). Cells were washed twice in saline and suspended in medium (RPMI 1640) supplemented with gentamicin (10 mg/mL), L-glutamine (10 nM) and pyruvate (10mM). The cells were counted in a Coulter counter (Beckman Coulter, Pasadena) and the number of was adjusted to 5 × 10^6^ cells/mL. A total of 5 × 10^5^ PBMCs were added in 100 ul to round-bottom 96-well plated (Greiner) and incubated with 100 uL of stimulus (heat-killed *Candida albicans* yeast of strain ATCC MYA-3573,UC 820, 1 ×10^6^/mL or RPMI 1640 as previously described(Matzaraki *et al*., 2021).

#### 1M-scBlood cohort

PBMCs from 120 volunteers were isolated and stimulated as previously described(Oelen *et al*., 2021). Briefly, we used Cell Preparation Tubes with sodium heparin (BD) to isolate PBMCs, which were cryopreserved until use in RPMI1640 containing 40% FCS and 10% DMSO. After thawing and a 1h resting period, unstimulated cells were washed twice in medium supplemented with 0.04% BSA and directly processed for scRNA-seq. On the other hand, for stimulation experiments, 5×105 cells were seeded in a nucleon sphere 96-well round bottom plate in 200 ul RPMI1640 (supplemented with 50 ug/mL gentamicin, 2 mM L-glutamine and 1 mM L-glutamine and 1mM pyruvate). The cells were stimulated with 1×10^6^ CFU/ml heat-killed *C. albicans* blastoconidia (strain ATCC MYA-3573, UC 820), 50 ug/ml heat-killed *M. tuberculosis* (strain H37Ra, Invivogen) or 1×10^7^ heat-killed *P. aeruginosa* (Invivogen) for 3h or 24h, at 37°C in a 5% CO2 incubator. After stimulations, cells were washed twice in medium supplemented with 0.04% BSA. Cells were then counted using a haemocytometer, and cell viability was assessed by Trypan Blue.

### Measurement of inflammatory proteins

Inflammatory protein concentrations were measured using Olink® proteomics platform. In fact, supernatants were collected after 24h of *Candida albicans* stimulation and submitted to Olink Proteomics for analysis using the inflammatory panel assay of 92 analytes. Olink data are presented as Normalized Protein eXpression values (NPX, based on log2 scale). Immunoassays utilized by Olink are based on the Proximity Extension Assay (PEA) technology(Assarsson *et al*., 2014), which makes use of oligonucleotide-labelled antibodies binding to their respective protein. When the two antibodies are brought in proximity, a DNA polymerase target sequence is formed, which is subsequently quantified by quantitative real-time polymerase chain reaction (qPCR).

### Preprocessing /filtering of protein data and normalization

Filtering of Olink generated data was restricted to only Proteins as the samples passed Olink internal quality control across all proteins. We excluded all proteins which failed to be quantified in at least 85% of the samples, meaning all proteins with more than 15% samples missingness (NPX value below the protein-specific limit of detection (LOD) value) were excluded from downstream analysis. The remaining Proteins with NPX values below the LOD were replaced with protein-specific LOD values.

Following per-protein cleaning, 42 and 35 proteins were available for the 1M-scBlodNL and 500FG cohorts respectively (**Figure S6A**). Out of these proteins, 26 were common between both cohorts.

The protein distributions on log2 scale were not normally distributed (**Figure S6B**). We applied rank-based inverse normal transformation as implemented in the GenABEL R package(Aulchenko *et al*., 2007), to transform the data to mimic Gaussian distribution(**Figure S6C**).

### SNP genotyping, quality control and imputation

The procedures for genotyping, genetic data filtering and genotype imputation of the 500FG cohort had been previously described(Li *et al*., 2016). Extracted DNA was genotyped using the commercially available SNP chip, Illumina HumanOmniExpressExome-8 v1.0. Following pre-imputation filtering steps for both markers and individuals, the remaining dataset SNP genotypes were imputed with GoNL as reference panel(Francioli *et al*., 2014).

For Lifelines Deep cohort, genotyping and imputation was performed as previously described(Ricaño-Ponce *et al*., 2016). Both the HumanCytoSNP-12 BeadChip and the ImmunoChip platforms (Illumina, San Diego, CA, USA) were used to genotype the isolated DNA. Independent markers quality control was performed for both platforms and subsequently merged into one dataset. After merging, genotype SNPs were imputed using IMPUTE2(Delaneau, Marchini and Zagury, 2012) against the GoNL reference panel.

### Correlation analysis

Pairwise Pearson correlation analysis using the ‘corrr” R package was performed on the normalized protein abundances after adjusting for age and sex. Based on Pearson’s correlations for each pair of proteins, co-expression protein networks were reconstructed (using significant correlation coefficient threshold of 0.7, (absolute(r) > 0.70), a cut-off denoting a very strong strength of association.

### Identification of pQTLs after Candida Albicans stimulation

The association analysis of genotype-phenotype correlation was carried out using two main approaches. In the first method, the univariate approach was performed using the linear regression in PLINK(Purcell *et al*., 2007). The pQTL analysis was conducted independently for both cohorts, that is one analysis for the 1M-scBloodNL and another for 500FG cohorts. In the second method, multivariate test of association based on canonical correlation analysis (CCA)(Ferreira and Purcell, 2009), was conducted. CCA extracts the linear combination of highly corrected traits that explain the largest possible amount of the covariation between genetic variants and all traits (Proteins in this case). To control for potential confounding factors, we adjusted for covariates such as age and sex on normalized protein abundances and, regressed the residuals of each protein and protein network on SNP genotypes.

### Meta-Analyses

Summary statistics from both primary studies were utilized to perform meta-analyses. Association results for both the 500FG and 1M-scBloodNL cohorts obtained from the univariate approach were combined using the weighted sum fixed-effect model as implemented in the METAL software program(Willer, Li and Abecasis, 2010). The multivariate approach implemented in PLINK does not compute beta estimates and standard errors. Therefore, the meta-analysis of the multivariate P-values was carried out using sum of z (Stouffer’s) method as implemented in the metap R package(Whitlock, 2005).

### Genetic colocalization analyses

Colocalization analysis of cell-type-specific *cis*-eQTL and pQTL signals was conducted using Bayesian colocalization method which is implemented in the coloc package in R(Giambartolomei *et al*., 2014). We retrieved the genome-wide cell-type-specific *cis*-eQTL summary statistics from a previously published study using the 1M-scBloodNL cohort(Oelen *et al*., 2021). Additionally, we performed trans-eQTL mapping only for the top pQTLs to ascertain their trans-eQTL effect. The eQTLs from this study were identified using PBMCs in an unstimulated condition as well as after 3h and 24h *in vitro*-stimulations with three different pathogen stimulations, namely *C. albicans, M. tuberculosis* and *P. aeruginosa. Cis*-eQTL was defined as SNP-gene distance of 100kb window and FDR < 0.05. However, for overlap comparison with pQTLs, we re-tested the relevant SNPs to ensure that SNPs outside the defined *cis*-region are included.

From our pQTL mapping results, we selected the index variants (SNPs with the smallest P value) for each protein and extracted all variants within a window size of 1Mb around the index pQTL variant for further analysis. Next, colocalization between Candidemia GWAS and pQTL signals were tested to ascertain whether the GWAS variants were colocalized with the index pQTL variant in a genomic region. For this analysis, we extracted summary statistics of genetic variants residing within 1Mb of the sentinel GWAS SNPs from previously published data(Jaeger *et al*., 2019). Given the lack of genome-wide significant hits reported in the GWAS study, a suggestive association threshold (P value < 1×10^−6^) was set to prioritize loci for further analysis. The default prior probability (1×10^−6^) that a random variant in the region is causal to both traits was applied. A posterior probability (PP4 >= 0.75) is considered as strong evidence of colocalization. LocusCompareR, being an R package was used for the visualization of results(Liu *et al*., 2019).

## Acknowledgments

The authors are most grateful to all volunteers from the 500FG and Lifelines DEEP cohorts for participation in the studies. Financial support. V.K was supported by a Hypathia tenure track grant RadboudUMC. MGN was supported by an ERC Advanced grant (#833247). M.W. was supported by a NWO Veni grant (#192.029). L.F. was supported by a NWO-VICI (#917.14.374) and an Oncode Investigator grant.

## Author contributions

V.K. & M.W. designed and supervised the study. M.G.N., M.W. & L.F. contributed with data generation and curation. C.K.B., R.O., & K.L. performed data analysis. C.K.B. performed pQTL mapping and other statistical analyses with critical input from V.K. C.K.B. drafted the manuscript & prepared the figures. All authors reviewed and edited the manuscript. All authors contributed to critical revisions and approved the final manuscript.

## Ethical approval and consent to participate

The 500FG cohort was approved by the Arnhem-Nijmegen Medical Ethical Committee (500FG: NL42561.091.12) and performed in accordance with the Declaration of Helsinki. All individuals gave written informed consent to donate venous blood for research.

The Lifelines DEEP study was approved by the ethics committee of the University Medical Center, Groningen, document number METC UMCG LLDEEP: M12.113965. All volunteers signed an informed consent from prior study of enrollment.

## Data and code availability

The cell-type specific eQTL summary statistics are available at the sc-eQTLGen website (https://eqtlgen.org/sc/datasets/1m-scbloodnl.html).

The summary-level association statistics of the meta-analyses results have been deposited at the EBI GWAS Catalogue under the accession number GCST90129614. In order not to compromise research volunteers’ privacy, individual level 500FG and 1M-scBloodNL genetic data can be acquired by researchers upon successful application using the data request from for the 500FG cohort (http://www.humanfunctionalgenomics.org/site/?page_id=16) and the 1M-scBloodNL cohort (https://eqtlgen.org/sc/datasets/1m-scbloodnl-dataset.html). The form will be reviewed by the Data Access Committee, who will grant access upon approval. The phenotypic data sets accompanying this manuscript are available on Dryad (https://doi.org/10.5061/dryad.k3j9kd5b6). No custom code was generated and all software that is central to the research findings have been previously reported and are referenced throughout the paper mostly in the methods section or as part of the figure legends.

